# Recently evolved combination of unique sulfatase and amidase genes enables bacterial degradation of the wastewater micropollutant acesulfame worldwide

**DOI:** 10.1101/2022.08.17.504299

**Authors:** Maria L. Bonatelli, Thore Rohwerder, Denny Popp, Yu Liu, Caglar Akay, Carolyn Schultz, Kuan-Po Liao, Chang Ding, Thorsten Reemtsma, Lorenz Adrian, Sabine Kleinsteuber

**Affiliations:** Department of Environmental Microbiology, Helmholtz Centre for Environmental Research – UFZ, Leipzig, Germany; Department of Environmental Biotechnology, Helmholtz Centre for Environmental Research – UFZ, Leipzig, Germany; Department of Analytical Chemistry, Helmholtz Centre for Environmental Research – UFZ, Leipzig, Germany; Institute of Analytical Chemistry, University of Leipzig, Leipzig, Germany; Chair of Geobiotechnology, Technische Universität Berlin, Berlin, Germany

## Abstract

Xenobiotics often challenge the principle of microbial infallibility. One example is acesulfame introduced in the 1980s as zero-calorie sweetener, which was recalcitrant in wastewater treatment plants until the early 2010s. Then, efficient removal has been reported with increasing frequency. By studying acesulfame metabolism in alphaproteobacterial degraders of the genera *Bosea* and *Chelatococcus*, we experimentally confirmed the previously postulated route of two subsequent hydrolysis steps via acetoacetamide-N-sulfonate (ANSA) to acetoacetate and sulfamate. Genome comparison of wildtype *Bosea* sp. 100-5 and a spontaneous acesulfame degradation-defective mutant revealed the involvement of two plasmid-borne gene clusters. The acesulfame-hydrolyzing sulfatase is strictly manganese-dependent and belongs to the metallo beta-lactamase family. In all degraders analyzed, it is encoded on a highly conserved gene cluster embedded in a composite transposon. The ANSA hydrolase, on the other hand, is an amidase signature domain enzyme encoded in another gene cluster showing variable length among degrading strains. Transposition of the sulfatase gene cluster between chromosome and plasmid explains how the two catabolic gene clusters recently combined for the degradation of acesulfame. Searching available genomes and metagenomes for the two hydrolases and associated genes indicates that the acesulfame plasmid evolved and spread worldwide in short time. While the sulfatase is unprecedented and unique for acesulfame degraders, the amidase occurs in different genetic environments and might have evolved for the degradation of other substrates. Evolution of the acesulfame degradation pathway might have been supported by the presence of structurally related natural and anthropogenic compounds, such as aminoacyl sulfamate ribonucleotide or sulfonamide antibiotics.

## INTRODUCTION

Acesulfame potassium is an artificial sweetener used in food and beverages, pharmaceuticals and personal care products. It was discovered in the late 1960s [1] and is a zero-calorie, high-potency sweetener that is 200 times sweeter than sucrose, with good storage and temperature stabilities [2]. It received its first approval of use in the UK in 1983 and has been used worldwide since the 1990s [2]. After human consumption, the acesulfame anion (ACE) is readily absorbed but not metabolized, and eventually excreted unchanged via the urinary tract. Consequently, ACE is present in domestic wastewater in a range between 10 and 100 μg L^−1^ [3] and can reach other aquatic environments [4]. Due to its hydrophilicity and persistence, ACE became an ideal anthropogenic wastewater tracer [3]. Recalcitrance of ACE in wastewater treatment plants (WWTPs) was reported first in 2009 [4], and its persistence in surface waters, groundwater and even tap water was observed in the USA [5, 6], Canada [7], China [8], Germany [9] and other countries in the early 2010s. However, reports on ACE biodegradation in WWTPs started to appear in the later 2010s. Two studies conducted in Australia [10] and China [11] revealed high percentages (92% and 85%, respectively) of ACE removal in WWTPs. When analyzing 13 WWTPs in Switzerland and Germany, Castronovo et al. [12] noticed 57% to 97% ACE removal and identified sulfamic acid/the sulfamate ion as the transformation product. Likewise, Kahl et al. [13] reported biodegradation of >85% in nine WWTPs in Germany and observed a seasonality of ACE removal with highest efficiency in summer and autumn. These results challenged the perception of ACE recalcitrance and clearly indicated biodegradation as the removal mechanism.

The reports on emerging ACE biodegradability in WWTPs inspired research on the microorganisms and metabolic pathways involved with this process. Kahl et al. [13] enriched ACE-degrading bacteria from sludge sampled from a treatment wetland in 2015. They used mineral medium with low concentrations of ACE (0.6 to 600 μM) as sole carbon source and confirmed its complete mineralization. The resulting community was dominated by *Alpha*- and *Gammaproteobacteria* (97 to 99%). Later, a pure culture identified as *Bosea* sp. (strain 3-1B) was isolated from this enriched community, and three more strains (*Bosea* sp. 100-5, *Chelatococcus* sp. 1g-2 and 1g-11) were isolated using fresh samples from the same wetland by direct enrichment on 0.5 to 5 mM ACE [14]. All strains degraded 100% of the ACE and formed sulfamate in stoichiometric amounts during batch cultivation. Additionally, acetoacetamide-N-sulfonate (ANSA) was detected transiently in culture supernatants, confirming the assumption of Castronovo et al. [12] who suggested ANSA as an intermediate of ACE degradation. Based on this finding, Kleinsteuber et al. [14] suggested a catabolic pathway of two subsequent hydrolysis steps via ANSA to acetoacetate (Fig. 1). More recently, Huang et al. [15] investigated ACE-degrading communities enriched from activated sludge and noticed that the most abundant metagenome-assembled genomes (MAGs) belonged to *Planctomycetes*, *Alpha*- and *Gammaproteobacteria*. These authors also isolated two ACE-mineralizing *Chelatococcus* strains (YT9 and HY11) and sequenced their genomes. They found a total of 812 protein-coding genes that were present in both strains but not in *Chelatococcus* type strains DSM 6462 and 101465, and speculated that the ACE degradation genes might be among them.

**Fig. 1.**
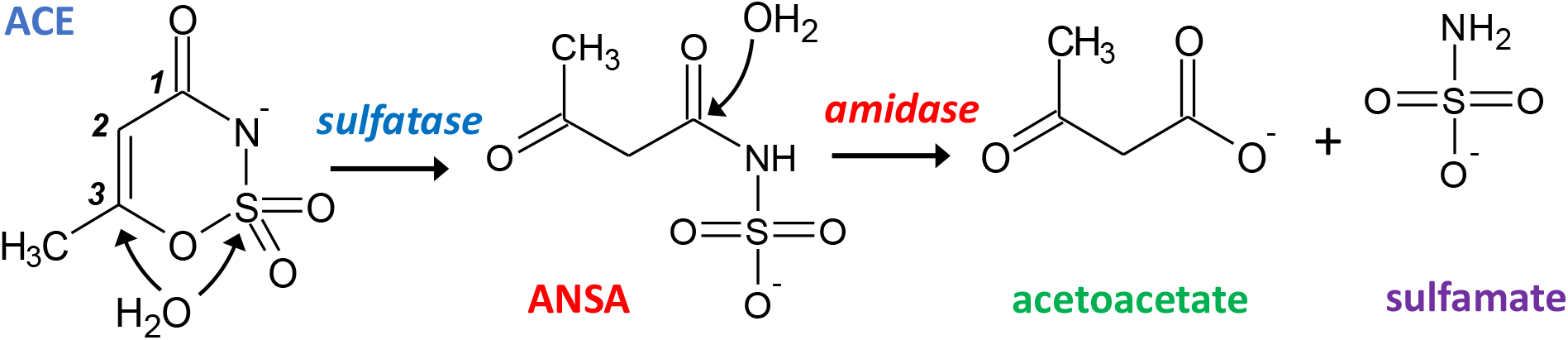
Proposed hydrolytic pathway for bacterial degradation of ACE. Initially, a sulfatase catalyzes the attack of either the sulfur or C3 atom of ACE resulting in the formation of ANSA. In a subsequent step, an amidase hydrolyzes the linear amide intermediate to acetoacetate and sulfamate.

Despite these advances, the biochemical and genetic background of this recently emerged catabolic trait as well as its evolutionary origin remained elusive. Here, we present insights into the ACE metabolism of our previously isolated ACE-degrading strains [14] and other strains isolated from WWTPs located in Germany. Based on comparative genomics and biochemical analysis, we identified the genes responsible for ACE degradation and surveyed their distribution in public sequence databases. Additionally, we discuss the mechanisms that might have established this pathway by combination of preexisting catabolic genes and facilitated its fast distribution by horizontal gene transfer.

## METHODS

### Isolation and cultivation of bacterial strains

*Bosea* sp. 3-1B was isolated from the ACE-degrading enrichment culture described by Kahl et al. [13] (inoculum sludge sampled in July 2015). *Bosea* sp. 100-5, *Chelatococcus* sp. 1g-2 and *Chelatococcus* sp. 1g-11 originated from the same treatment wetland sampled in February 2019. The strains were isolated by plating on R2A agar after enrichment in DSMZ 462 mineral medium with 0.5 or 5 mM ACE as sole carbon source [14]. *Chelatococcus asaccharovorans* WSA4-1, *Chelatococcus* sp. WSC3-1, *Chelatococcus asaccharovorans* WSD1-1 and *Chelatococcus* sp. WSG2-a were isolated from enrichment cultures in DSMZ 461 mineral medium with 5 mM ACE inoculated with activated sludge from four WWTPs in Germany (sampled in 2020, see Table 1). The strains were identified by Sanger sequencing of 16S rRNA genes and maintained on DSMZ 461 agar with 5 mM ACE incubated at 30°C. Liquid cultures to study growth and degradation kinetics or for enzyme assays and DNA extraction were set up in DSMZ 461 or DSMZ 462 medium with the respective carbon source (2.5 to 10 mM ACE, 10 mM 3-hydroxybutyrate or 10 mM succinate).

**Table 1.** Information on the bacterial genomes analyzed in this study.

| Strain | Sampling site/Isolation origin | Sampling date | Localization of metabolic clusters 1 and 2 | Contig size of the metabolic clusters (bp) | Assembly accession | Reference |
| --- | --- | --- | --- | --- | --- | --- |
| *Bosea* sp. 100-5 | Treatment wetland Langenreichenbach, Mockrehna, Germany (51°30'N 12°53'E) | 2019 | Contig 4 | 40 931 | GCA_930633465 | [14] |
| *Bosea* sp. 100-5 Mut1 | Spontaneous mutant of *Bosea* sp. 100-5 | isolated in 2021 | Contig 56 (only cluster 2) | 9 696 (only cluster 2) | GCA_946047605 | This work |
| *Bosea* sp. 3-1B | Treatment wetland Langenreichenbach, Mockrehna, Germany (51°30'N 12°53'E) | 2015 | Contig 3 | 43 141 | GCA_930633495 | [14] |
| *Chelatococcus* sp. 1g-2 | Treatment wetland Langenreichenbach, Mockrehna, Germany (51°30'N 12°53'E) | 2019 | Contig 3 | 41 507 | GCA_930633515 | [14] |
| *Chelatococcus* sp. 1g-11 | Treatment wetland Langenreichenbach, Mockrehna, Germany (51°30'N 12°53'E) | 2019 | Contig 6 | 45 665 | GCA_930633505 | [14] |
| *Chelatococcus asaccharovorans* WSA4-1 | WWTP Markranstädt, Germany (51°18'N 12°12'E) | 2020 | Contig 5 | 45 942 | GCA_930633525 | This work |
| *Chelatococcus* sp. WSC3-1 | WWTP Rosental, Leipzig, Germany (51°21'N 12°20'E) | 2020 | Contig 2 | 45 505 | GCA_930633455 | This work |
| *Chelatococcus asaccharovorans* WSD1-1 | WWTP Markkleeberg, Germany (51°17'N 12°21'E) | 2020 | Contig 6 | 41 507 | GCA_930633485 | This work |
| *Chelatococcus* sp. WSG2-a | WWTP Wiedemar, Germany (51°28'N 12°11'E) | 2020 | Contig 1 | 45 506 | GCA_930633475 | This work |
| *Chelatococcus* sp. YT9 | WWTP Shatin, Hong Kong, China | 2018 | Contig 1 (cluster 1)<br>Contig 5 (cluster 2) | 4 368 288 (cluster 1)<br>16 496 (cluster 2) | GCA_018398315 | [15] |
| *Chelatococcus* sp. HY11 | WWTP Shatin, Hong Kong, China | 2018 | Contig 6 (only cluster 2) | 18 060 (only cluster 2) | GCA_018398335 | [15] |

To select an ACE degradation-defective mutant, *Bosea* sp. 100-5 was grown in R2A medium for twelve transfers and plated on R2A agar in dilution series to obtain single colonies. Colonies were isolated and screened for growth in liquid culture on either ACE or succinate as sole carbon source. One isolate that grew on succinate but not on ACE was identified and designated as *Bosea* sp. 100-5 Mut1.

### Genome sequencing

Genomic DNA was extracted from ACE-grown cells (DSMZ medium 461 with 5 mM ACE) with the NucleoSpin^®^ Microbial DNA kit (Macherey-Nagel, Düren, Germany) according to the manual and using wide-bore tips. DNA concentration was measured with the Qubit dsDNA BR Assay Kit (Thermo Fisher Scientific, Rockford, IL, USA); DNA integrity was checked by gel electrophoresis.

Genomes of all strains except the mutant *Bosea* sp. 100-5 Mut1 were sequenced using a long-read and short-read approach. Long-read sequencing was done on the MinION MK1b platform (Oxford Nanopore Technologies, Oxford, UK) as previously described [16] with the exception that Guppy version 4.0.15 was used for basecalling. Nanopore read data was downsampled per isolate to about 150x coverage using filtlong (https://github.com/rrwick/Filtlong) with a priority on read length (--length_weight 10). Short-read sequencing of strain 3-1B was conducted on the Illumina MiSeq platform (Illumina, San Diego, CA, USA) using the NEBNext Ultra II FS DNA library prep kit (New England Biolabs, Frankfurt, Germany) to prepare a 2×300 bp library. Short-read sequencing of the other strains was performed by Genewiz GmbH (Leipzig, Germany) on the Illumina NovaSeq platform (2×150 bp). Short and long reads were assembled with a hybrid approach using Unicycler v0.4.9 [17] with default parameters.

Genome sequencing of *Bosea* sp. 100-5 Mut1 was conducted on the Illumina NextSeq 2000 platform (2×150 bp) by StarSeq GmbH (Mainz, Germany). After assembly with SPAdes version 3.15.2 [18], contigs smaller than 500 bp were excluded from the assembly. Platanus version 1.2.1 [19] was used for scaffolding and Mauve [20] for reordering the contigs using the wildtype genome of *Bosea* sp. 100-5 as reference.

### Genome annotation and comparative genomics

The MicroScope automatic annotation pipeline was used for the prediction and functional annotation of coding sequences (CDS) [21]. GTDB-tk [22] and CheckM [23] were used for taxonomic classification and estimating genome completeness and contamination, respectively. The annotated genomes were submitted to the NCBI under the accession numbers given in Table 1; genome metrics is provided in Table S1. To compare the genome of *Bosea* sp. 100-5 with its mutant *Bosea* sp. 100-5 Mut1, we used Geneious 10.0.9 [24], Mauve and Proksee [25]. Additionally, the Synteny Statistics tool of MicroScope was used to explore the similarity between the two genomes. Gene clusters of interest and surrounding genes were investigated in all genomes with cblaster version 1.3.11 [26]. Besides the genomes sequenced in this study, we included the genomes of *Chelatococcus* sp. strains YT9 (JAHBRW01) and HY11 (JAHBRX01) as well as the MAG sequences of the corresponding BioProject (PRJNA725625) [15]. Briefly, protein sequences of >100 amino acids were searched against a database built from genomes annotated with MicroScope applying default parameters, except for minimal identity of 90%. For the genomes that did not present the gene clusters, tblastn was used to manually search for the individual CDS BOSEA1005_40015.

To look for genes related to BOSEA1005_40015 in public databases, we searched the NCBI (https://www.ncbi.nlm.nih.gov/taxonomy) and the JGI (https://img.jgi.doe.gov/cgi-bin/m/main.cgi) databases (both August 2022). See Supplementary information for further details. For datasets presenting hits for the query sequence BOSEA1005_40015, we searched for other elements of the *Bosea* sp. 100-5 plasmid and used Geneious 10.0.9 and Proksee to compare them. Predicted proteins with ambiguous annotation were searched for functional sites with the NCBI Conserved Domain tool [27] and InterPro [28].

### Biochemical analyses

Crude extracts from bacterial cells were prepared in lysis buffer (10 mM Tris-HCl with 10% glycerol, pH 7.8) applying French press or disruption in a mixer mill with glass beads [29]. Extracts were obtained by centrifugation at 20 000 x g and 4°C for 20 min. For removal of low molecular weight compounds interfering with HPLC analysis, protein extracts were purified with 10 kDa Amicon filters (Merck, Darmstadt, Germany). Protein concentration was determined with Pierce BCA Protein Assay Kit (Thermo Fisher Scientific) or Bradford reagent (AppliChem, Darmstadt, Germany) using bovine serum albumin as standard. For degradation assays and as analytical standards, ACE potassium salt (99% pure, Merck) and acetoacetate lithium salt (95% pure; abcr GmbH, Karlsruhe, Germany) were used. As ANSA is not commercially available, it was prepared employing the ACE-hydrolyzing activity of *Bosea* sp. 100-5. To separate ACE and ANSA hydrolase activities, size exclusion chromatography was used with protein crude extracts of ACE-grown cells, a Superdex 200 10/300 column (Cytiva, Marlborough, MA, USA) and 100 mM ammonium hydrogen carbonate buffer pH 7.82 as eluent at 0.5 ml min^−1^. The ACE hydrolase eluted with proteins of around 70 kDa, while ANSA-hydrolytic activity was found in fractions representing 150 to 240 kDa. The 70-kDa fraction was directly used to prepare up to 20 mM ANSA at 2-mL scale. The conversion was stoichiometric enabling a calibration for ANSA. Proteins were removed by ultrafiltration (3 kDa Vivaspin; Sartorius, Göttingen, Germany). ACE and ANSA hydrolysis activities were quantified in discontinuous HPLC-based assays at 30°C in lysis buffer. Samples were diluted in stop buffer (10 mM malonate, pH 4.0, 60°C) prior to HPLC analysis.

For heterologous expression, the genes BOSEA1005_40016 and _40030 were synthesized and cloned into pET-28a(+)-TEV via *Nde*I/*Bam*HI sites (GenScript, Oxford, UK). Transformed *E. coli* Lemo21 (DE3) (New England Biolabs) was incubated in lysogeny broth with 30 ppm chloramphenicol, 50 ppm kanamycin and 0.1 mM rhamnose at 30°C until an optical density (600 nm) of 0.5. Then, 0.4 mM isopropyl β-d-1-thiogalactopyranoside and 2 % (v/v) ethanol were added and incubation proceeded at 17°C for 20 h.

### Analytics

ACE and its metabolites ANSA and acetoacetate were quantified by HPLC with an UV/VIS detector employing a Nucleosil 100-5 C18 HD column (240/3 mm, Macherey-Nagel) and an eluent at 0.5 ml min^−1^ containing 100 mM NaH2PO4 and 10 mM tetrabutylammoniumhydrogensulfate, pH 4.5, and an acetonitrile gradient (% v/v): 2 min 7.5, linear increase to 27.5 in 2 min, 6 min 27.5, linear decrease to 7.5 in 2 min, 12 min 7.5. ACE and ANSA peaks were analyzed at 260 nm, acetoacetate at 274 nm.

Identity of enzymatically prepared ANSA was validated by LC-MS operated with electrospray ionization in negative polarity and equipped with a Zorbax Eclipse Plus Rapid Resolution HT-C18 column (100 x 3 mm, 1.8 μm) at 30°C. A binary mobile phase at 0.4 ml min^−1^ consisted of 0.2% formic acid and a methanol gradient: 1 min equilibration at 10%, 2 min linear increase to 55%, 7 min linear increase to 95%, 2 min hold at 95%, 0.5 min decrease to 10% and 3.5 min hold at 10%. Further details are given in the Supplementary information.

## RESULTS

### Enzymatic ACE hydrolysis in *Bosea* sp. 100-5 proceeds via ANSA to acetoacetate and strongly depends on Mn^2+^ ions

ACE degradation to acetate via ANSA (Fig. 1) was experimentally confirmed with strain *Bosea* sp. 100-5. Protein crude extracts from ACE-grown cells readily degraded ACE to acetoacetate (Fig. 2a) with transient accumulation of ANSA (Fig. 2b). Enzyme activity did not depend on addition of any cosubstrates, such as NADH or ATP, confirming that ACE degradation to acetoacetate is a two-step hydrolytic reaction as already postulated [14]. Moreover, growth on ACE as sole source of carbon and energy was strongly dependent on supplementation with Mn^2+^ ions. In mineral medium with excess Mn^2+^, ACE degradation allowed exponential growth with doubling times of about 19 h (Fig. 2c). Growth and ACE turnover was substantially reduced in Mn^2+^-limited medium (Fig. 2d). Supplementation with other divalent metals, such as Zn^2+^, Fe^2+^, Cu^2+^ or Co^2+^, did not reverse this effect. However, *Bosea* sp. 100-5 grew well in Mn^2+^-limited medium on alternative carbon sources, such as succinate or 3-hydroxybutyrate. The latter substrate is likely metabolized through the same route employed for the carbon skeleton of ACE, i.e. via acetoacetate, acetoacetyl-CoA and acetyl-CoA. Hence, the observed metal dependence is specific for growth on ACE, suggesting that one or both hydrolytic steps leading to the intermediate acetoacetate are Mn^2+^-dependent.

**Fig. 2.**
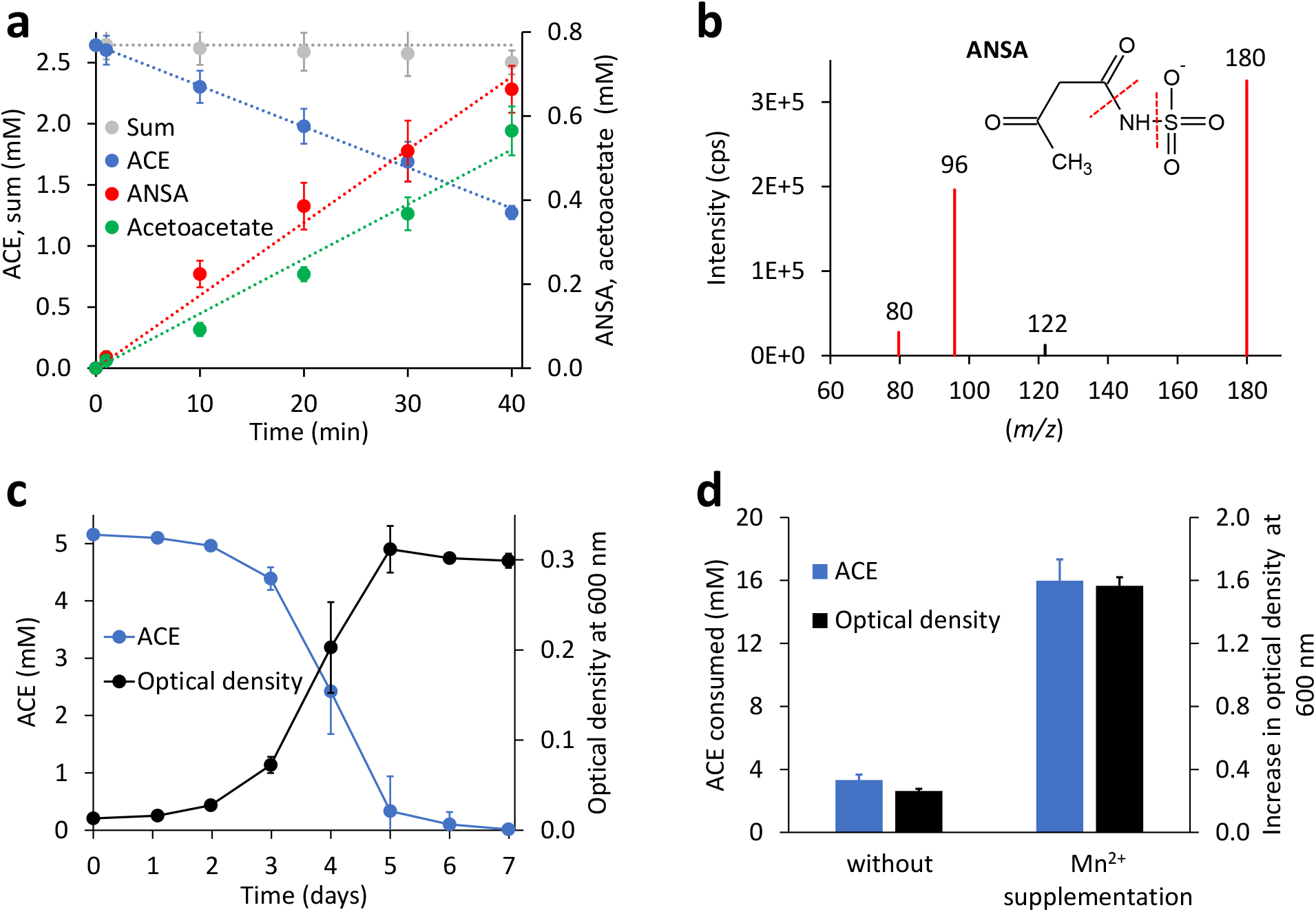
Physiological and biochemical characteristics of bacterial ACE degradation. **a** ACE degradation assay at 30°C with crude protein extract (470 μg mL^−1^ total protein) obtained from *Bosea* sp. 100-5 cells grown on ACE as sole source of carbon and energy. **b** Identification of ANSA (anion *m/z* = 180) as product of enzymatic ACE hydrolysis. Enhanced product ion spectrum obtained by LC-MS/MS analysis with electrospray ionization in negative polarity. Formation of characteristic products ([SO_3_]^−^ with *m/z* = 80 and [H_2_NSO_3_]^−^ with *m/z* = 96) is indicated. **c** Batch growth of *Bosea* sp. 100-5 on 5 mM ACE at 30°C in Mn^2+^-rich mineral medium (66.7 μM Mn^2+^) allowing doubling times as low as 19 h. **d** Metal dependence of fed-batch growth and degradation of ACE in *Bosea* sp. 100-5 cultivated in Mn^2+^-limited mineral medium (0.15 μM Mn^2+^) without or with Mn^2+^ supplementation (3.6 μM Mn^2+^). Cultures were incubated for 5 days at 30°C with a total amount of 20 mM ACE. Values given represent mean and SD of at least five independent experiments.

### Two plasmid-borne gene clusters are involved in ACE degradation

The ACE degradation trait seemed to be unstable in *Bosea* sp. 100-5 and was lost after prolonged cultivation in complex medium without ACE. The obtained mutant strain *Bosea* sp. 100-5 Mut1 could neither grow on ACE nor consume it in the presence of the alternative carbon source 3-hydroxybutyrate. Moreover, crude extracts did not transform ACE, suggesting that not merely a substrate uptake system but directly the initial hydrolytic step was affected in the mutant.

Mapping the genome of *Bosea* sp. 100-5 Mut1 on the corresponding wildtype genome revealed that only 2.5% of the nucleotides are missing in the mutant genome and 98.15% of the CDS are conserved among the two genomes. The wildtype genome comprises a contig of 40 931 bp that was annotated as a plasmid (Fig. 3). A gap of 26 252 bp was found in the corresponding region of the mutant genome (Fig. 3 and S1). This deletion comprises a putative metabolic gene cluster of six CDS encoding two hydrolases and four transport proteins, flanked by several CDS typically associated with a composite transposon, such as genes encoding transposases, integrases and transposase-assisting ATPases (Fig. 3, Table 2).

**Fig. 3.**
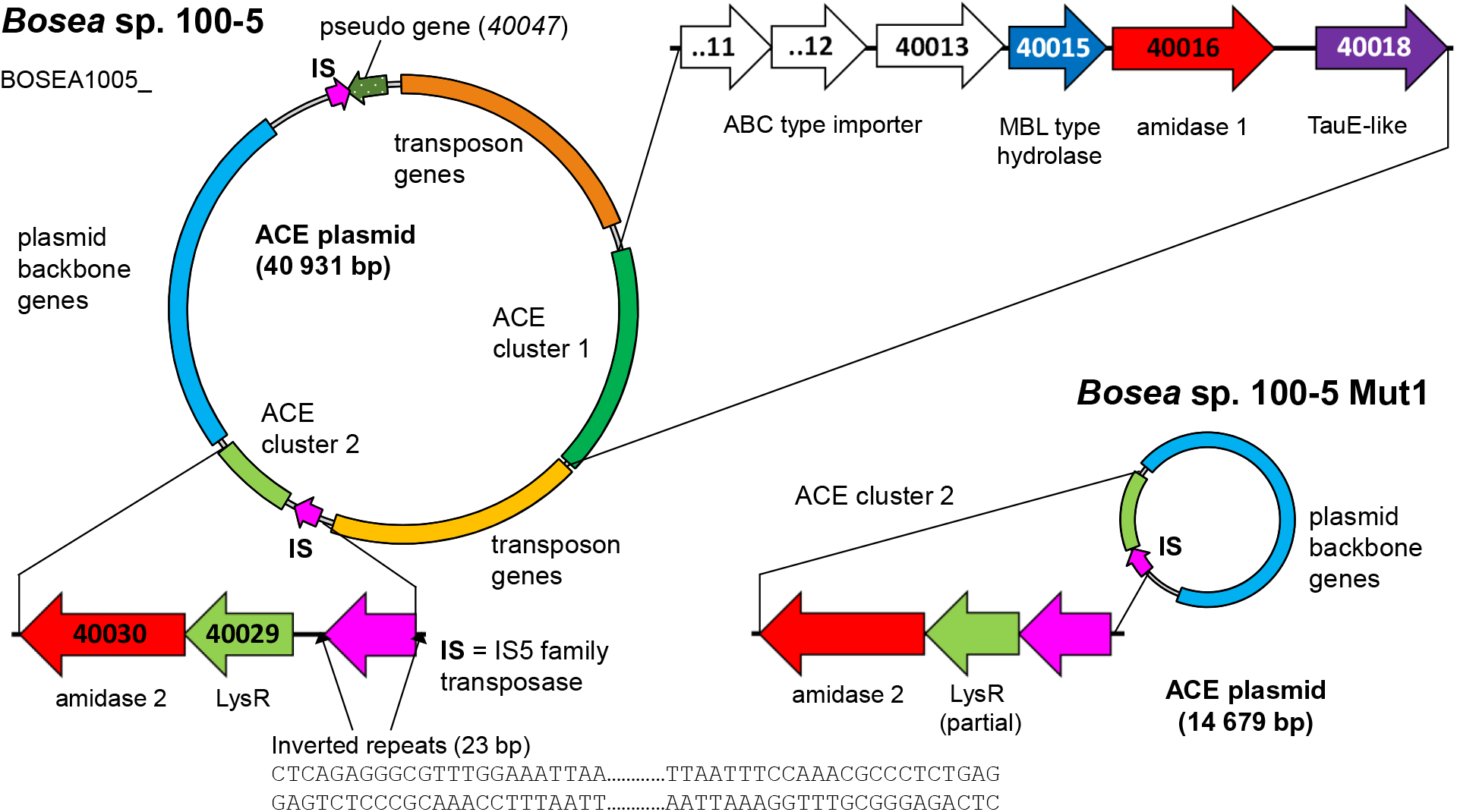
Deletion of the 22.5-kb composite transposon and additional genes from the 40.9-kb ACE plasmid found in *Bosea* sp. 100-5. Within the transposon, the wildtype plasmid bears a gene cluster (ACE cluster 1) encoding two hydrolyses and four transport proteins. Additionally, a second gene cluster encoding another hydrolase (ACE cluster 2), plasmid backbone genes and two copies of an IS5 family transposase (IS) are present. In the mutant strain *Bosea* sp. 100-5 Mut1, a 14.7-kb plasmid containing the backbone genes and ACE cluster 2 remained. The plasmid was manually reconstructed from contigs 56, 57 and 70 of the Mut1 genome sequence (GCA_946047605). Numbers in genes refer to locus tags in *Bosea* sp. 100-5 (prefix BOSEA1005_).

**Table 2.**
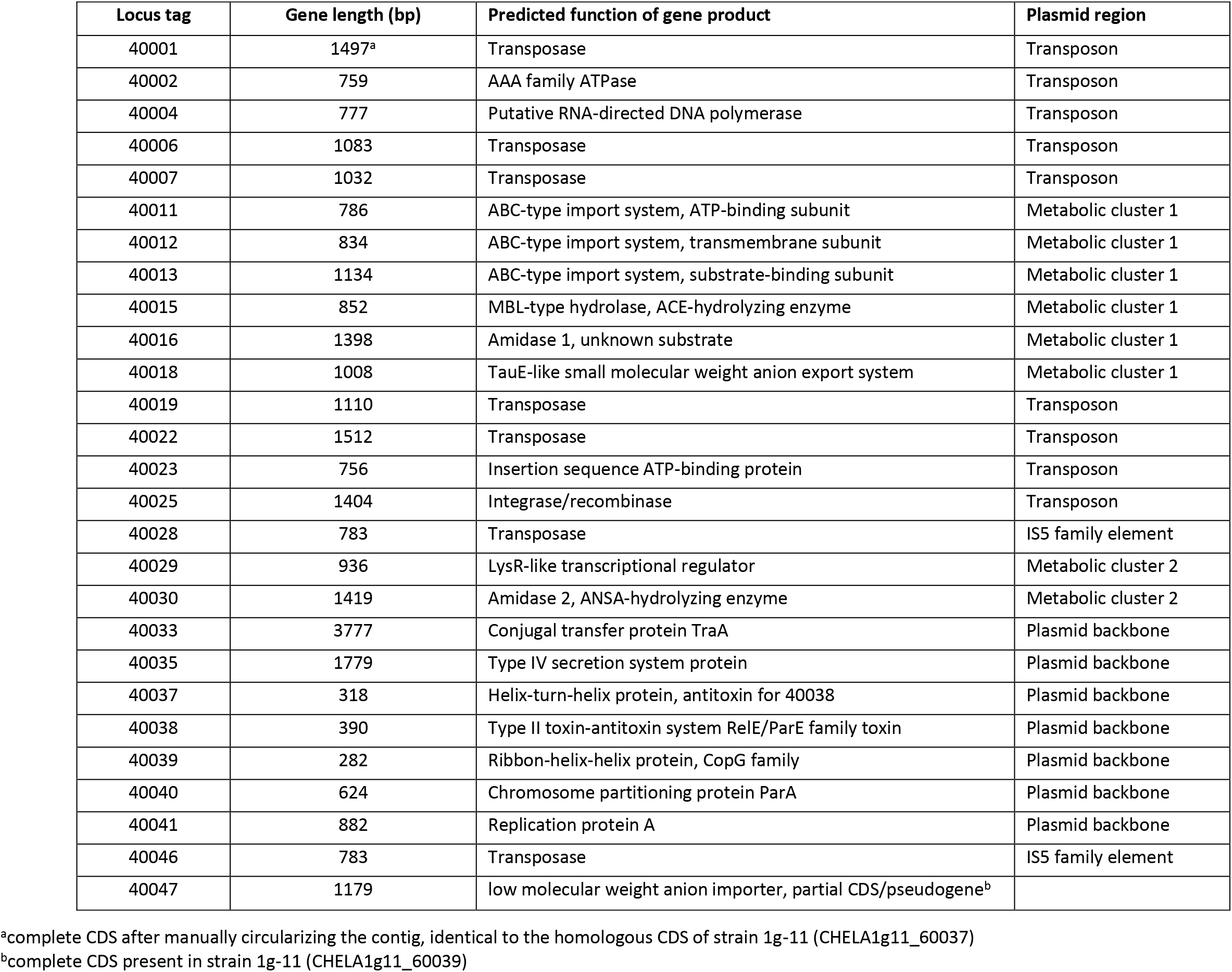
CDS of the ACE plasmid from *Bosea* sp. 100-5 with predicted functions (locus tag prefix BOSEA1005_).

Three CDS of the identified cluster (BOSEA1005_40011, 40012 and 40013) encode the essential components of an ATP-binding cassette (ABC) transport system potentially involved in ACE uptake (see also Supplementary information for functional assignment and sequence comparison of predicted proteins). The two hydrolases encoded by BOSEA1005_40015 and 40016 belong to the metallo beta-lactamase (MBL)-type and amidase signature sequence enzyme families, respectively. BOSEA1005_40018 encodes a protein related to the TauE-like anion transport system, which might be responsible for the removal of the ANSA hydrolysis product sulfamate.

Catalytic activity of MBL-type hydrolases depends on divalent metal ions coordinated by highly conserved active site His and Asp residues [30]. Considering the observed Mn^2+^ dependence of ACE degradation in *Bosea* sp. 100-5 (Fig. 2d), the MBL-type hydrolase (BOSEA1005_40015) is the most likely candidate for the initial attack on ACE. In contrast, enzymes with amidase signature sequence do not show any specific metal dependence but possess a unique Ser Ser Lys catalytic triad for the nucleophilic attack of the carbonyl carbon found in amide groups [31]. Consequently, it was tempting to assign the hydrolytic activity for ANSA degradation to BOSEA1005_40016. However, directly downstream of the composite transposon, a second metabolic gene cluster was found (Fig. 3) encoding a LysR-like transcriptional regulator (BOSEA1005_40029) and a second enzyme of the amidase signature sequence family (BOSEAS1005_40030, 39% identity with BOSEA1005_40016). The latter genes are still present in the mutant plasmid (Fig. 3). In crude extracts from 3-hydroxybutyrate-grown Mut1 cells, ANSA was readily converted to acetoacetate, ruling out BOSEA1005_40016 as the primary ANSA hydrolase. Moreover, crude extracts of *E. coli* after heterologous expression of BOSEA1005_40030 showed high ANSA-hydrolyzing activity (Fig. 4a), while heterologous expression of BOSEA1005_40016 did not confirm ANSA hydrolysis. In conclusion, the two-step hydrolysis of ACE to acetoacetate and sulfamate (Fig. 4b) is encoded by two different gene clusters located on one plasmid in *Bosea* sp. 100-5 (Fig. 3).

**Fig. 4.**
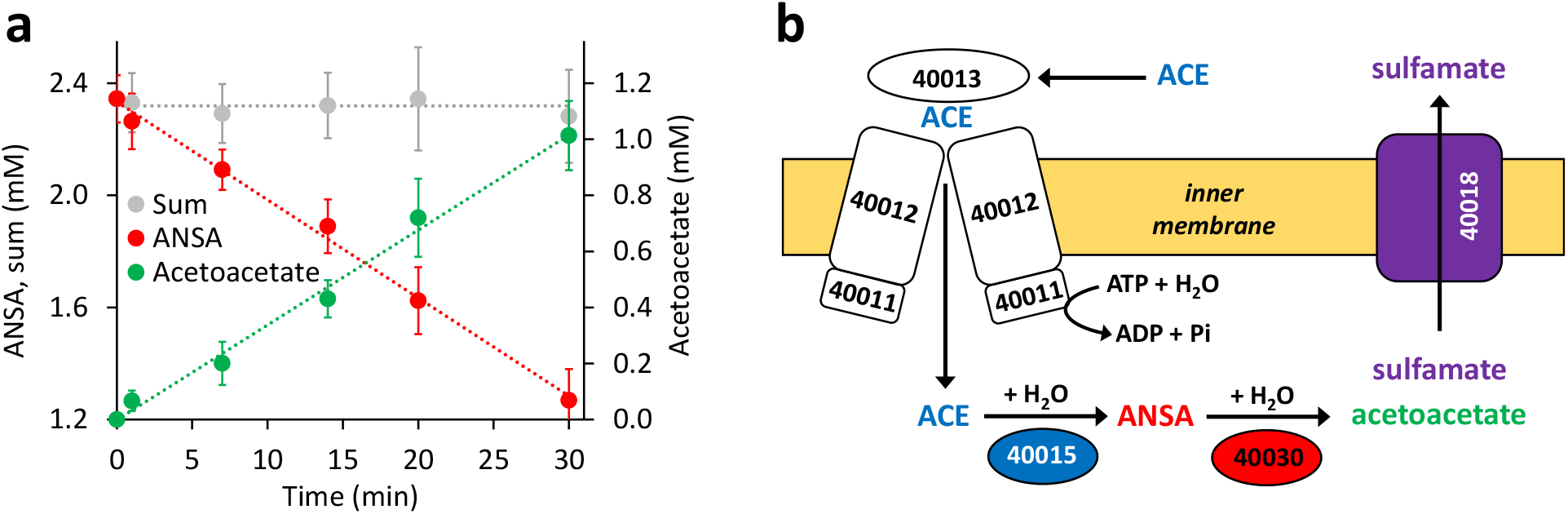
Both hydrolase-encoding gene clusters found on the 40.9-kb ACE plasmid of the wildtype *Bosea* sp. 100-5 are involved in ACE degradation. **a** Stoichiometric conversion of ANSA to acetoacetate in crude extract (74 μg mL^−1^ total protein) from *E. coli* Lemo21 (DE3) after expressing the BOSEA1005_40030 gene. Values given represent mean and SD of at least five independent experiments. **b** Proposed scheme of the two-step hydrolysis of ACE catalyzed by cytoplasmic MBL-type hydrolase BOSEA1005_40015 (from ACE cluster 1) and amidase BOSEA1005_40030 (from ACE cluster 2). Likely, ACE and its final hydrolysis product sulfamate are taken up and removed, respectively, by the transport systems encoded in ACE cluster 1 (see Fig. 3), but both assumptions need experimental support. Numbers shown refer to locus tags in *Bosea* sp. 100-5 (prefix BOSEA1005_).

### The plasmid-borne metabolic genes found in *Bosea* sp. 100-5 are highly conserved among ACE degraders

In addition to the previously described strains *Bosea* sp. 100-5, *Bosea* sp. 3-1B, *Chelatococcus* sp. 1g-11 and *Chelatococcus* sp. 1g-2 [14], we isolated several other ACE-degrading *Chelatococcus* sp. strains from samples collected in WWTPs from different locations in Germany (Table 1) and sequenced their genomes (Table S1). In agreement with their proposed role in ACE degradation, the two gene clusters were found in all genomes on plasmid-like contigs ranging from 41 507 to 45 942 bp (Table 1). While ACE cluster 1 (sulfatase cluster) together with its flanking transposon-related CDS is highly conserved (>99% similarity) among all strains, ACE cluster 2 (amidase cluster) is variable in size (two to four CDS) but always contains the conserved CDS for the ANSA amidase and the LysR-like transcriptional regulator. The plasmid backbones are also similar among the compared genomes, indicating that a specific type of conjugative plasmid is linked with the ACE degradation trait in all these strains. The ACE plasmid of *Chelatococcus* sp. 1g-11 is exemplarily shown in Fig. 5.

**Fig. 5.**
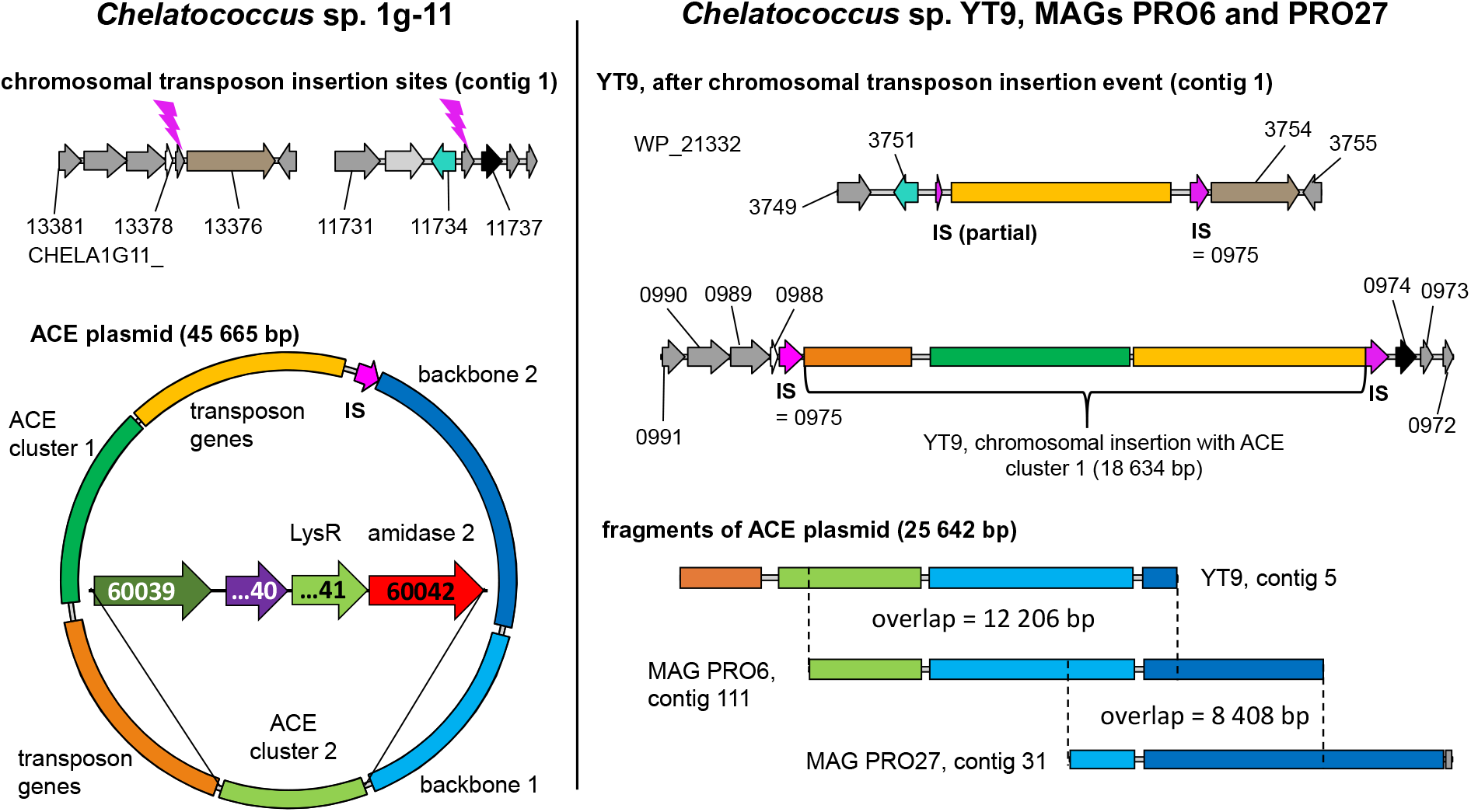
Tracing back the transposition event resulting in the insertion of the transposon bearing the ACE cluster 1 into the chromosome of strain *Chelatococcus* sp. YT9. Left panel: *Chelatococcus* sp. 1g-11, chromosomal insertion sites (GCA_930633515, contig 1) for the IS5 family transposase (IS) encoded on the 45.7-kb ACE plasmid (GCA_930633515, contig 6) that also bears the ACE sulfatase and ANSA amidase genes on ACE clusters 1 and 2, respectively. Compared to the corresponding plasmid found in wildtype *Bosea* sp. 100-5 (Fig. 3), only a part of the backbone genes is shared (“backbone 1”), whereas other genes are only present in strain 1g-11 (“backbone 2”). Right panel: *Chelatococcus* sp. YT9, chromosomal sites affected by the transposon insertion event (NZ_JAHBRW010000001.1). At one site, the transposon with ACE cluster 1 is integrated. From the ACE plasmid itself, however, only a small fragment is present in the genome of strain YT9 (NZ_JAHBRW010000005.1). Additional fragments (JAHBYS010000111.1, JAHBYU010000031.1) can be found in MAGs obtained from the ACE-degrading enrichment cultures from which also strain YT9 was isolated. Numbers marking genes refer to locus tags (with prefix as indicated). Genes with the same color refer to identical or closely related sequence (≥80% identity, in most cases >99%).

### Genome rearrangements in *Bosea* and *Chelatococcus* strains illustrate genetic plasticity of the ACE degradation trait

In the mutant strain *Bosea* sp. 100-5 Mut1, the composite transposon comprising ACE cluster 1 has been completely deleted from the plasmid (Fig. 3). A similar situation was found in the published genome of the ACE-degrading strain *Chelatococcus* sp. HY11 isolated from a WWTP in Hong Kong [15]. Here, ACE cluster 1 is missing, like most of the other CDS of the composite transposon. However, some genes corresponding to the plasmid backbone are located on a short contig of the HY11 genome (JAHBRX010000006.1). It is worth mentioning that the plasmid backbone gene for the type IV secretion system protein (Table 2) is disrupted by a Tn3 family transposase gene (MBS7743782.1), indicating some recombination events in this genomic region of strain HY11. Likewise, various fragments of the rudimentary plasmid, lacking most of the 26-kb region associated with ACE hydrolysis, are present in the MAG sequences PRO6 and PRO27 (Fig. 5). The corresponding metagenome was obtained from samples of ACE-degrading consortia seeded with activated sludge from the WWTP in Hong Kong, from which also the ACE degraders *Chelatococcus* sp. HY11 and YT9 were isolated [15].

While the genome sequences of *Bosea* sp. 100-5 Mut1 and likely also of *Chelatococcus* sp. HY11 document the loss of ACE cluster 1, the genome of *Chelatococcus* sp. YT9 gives evidence for the insertion of the composite transposon into the chromosome. Here, ACE cluster 1 is flanked by almost all transposon components found in the 26 kb region of *Bosea* sp. 100-5 (Fig. 5). Due to the insertion event, identical copies of an IS5 family transposase gene frame the whole gene cluster (WP_213320975.1; 100% identical with BOSEA1005_40028 and BOSEA1005_40046). As the insertion sites are well conserved in *Chelatococcus* sp. 1g-11, the event can be reconstructed (Fig. 5). Likely, the mobile element was originally located on a plasmid as the ones found in our ACE-degrading isolates (Table 1). Accordingly, short contigs of strain YT9 and MAGs PRO6 and PRO27 harbor parts of the transposon genes, ACE cluster 2 and some of the backbone genes of the ACE plasmid found in *Chelatococcus* sp. 1g-11.

### Putative ACE degradation gene clusters in public sequence databases

The ACE sulfatase (BOSEA1005_40015) appears to be a unique feature of the pathway. While it is ≥99% conserved among all genome-sequenced ACE degraders thus far (Table 1), only very distantly related sequences (with <30% identity at ≥80% coverage) were found in other published genomes (bacterial isolates and MAGs). Therefore, we surveyed public metagenomes and metatranscriptomes from wastewater environments and found the ACE sulfatase gene in sequence datasets from activated sludge sampled in Austria, China, Germany, Netherlands, Taiwan and USA (Fig. 6). As the samples were collected between 2015 and 2018, the first record of the ACE sulfatase gene in 2015 can be traced back to the Klosterneuburg WWTP in Austria (Table S2).

**Fig. 6.**
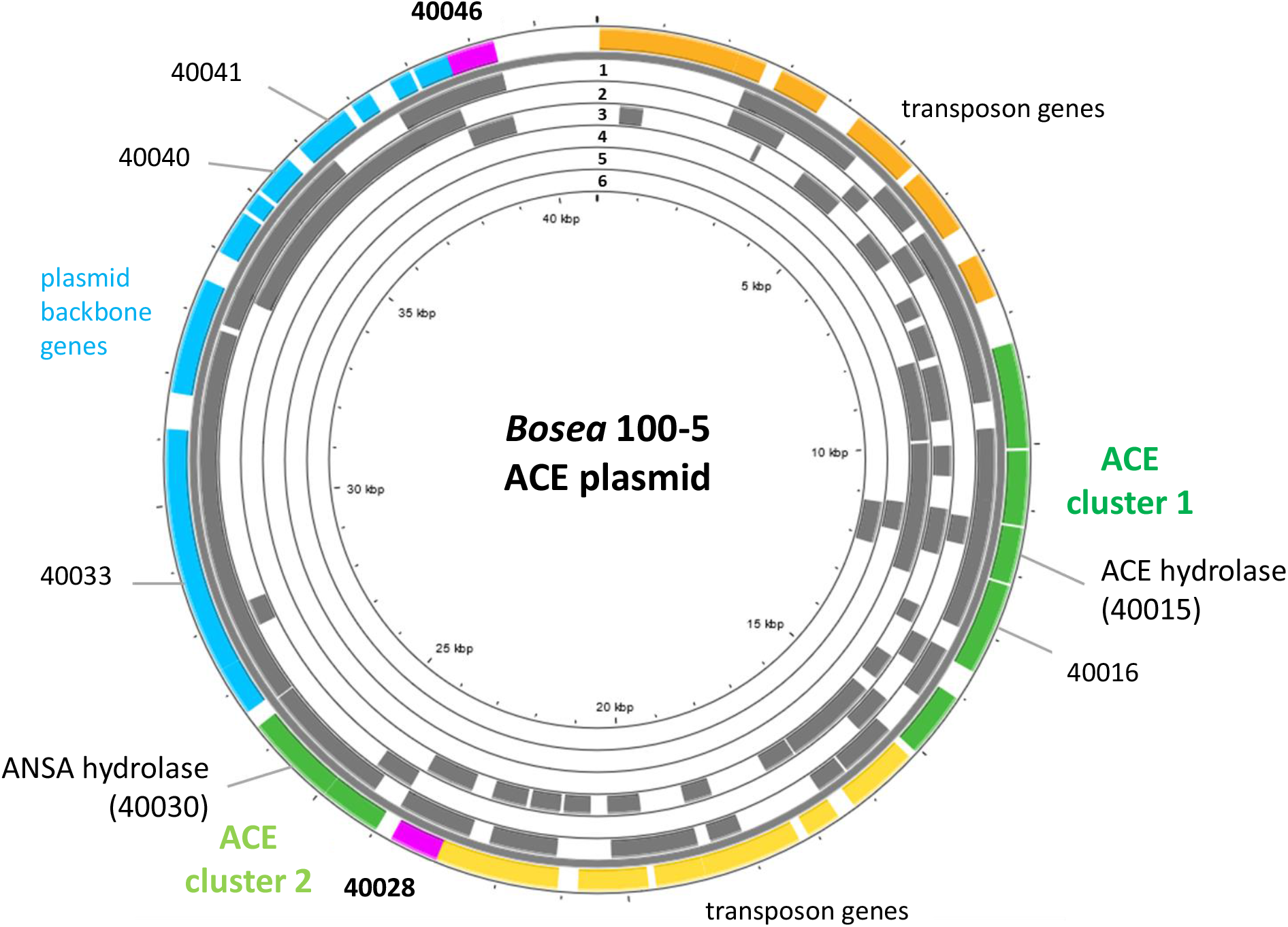
Mapping of the ACE plasmid genes on metagenome and metatranscriptome datasets associated with wastewater, bioreactors, and WWTPs. The outer colored ring shows the genes on the *Bosea* sp. 100-5 ACE plasmid (color code as in Fig. 3; genes highlighted, besides 40015 and 40030, encode an amidase of unknown function, 40016, the IS5 family transposase, 40028 and 40046, conjugal transfer protein TraA, 40033, chromosome partitioning protein ParA, 40040, and replication protein A, 40041, locus tag prefix is BOSEA1005_). For complete annotation, see Table 2. Grey areas show hits within the following metagenome and metatranscriptome datasets: 1 = Anaerobic bioreactor from Hunan University, China; 2 = Linkou WWTP, Taiwan; 3 = Wenshan WWTP, Taiwan; 4 = Weurt WWTP, The Netherlands; 5 = Klosterneuburg WWTP, Austria; 6 = WWTP in Virginia, USA. For complete matches from datasets, see Table S2. The figure was generated with Proksee.

In some datasets that presented the ACE sulfatase gene, other elements of the ACE plasmid from *Bosea* sp. 100-5 were also found. In the metatranscriptome from the Weurt WWTP in Nijmegen (Netherlands), genes of ACE cluster 1 corresponding to BOSEA_40011 to _40016 were detected, and the metagenome from the Wenshan WWTP (Taiwan) was shown to contain the complete ACE cluster 1. In the latter dataset, we also found contigs related to plasmid backbone and transposon genes of the *Bosea* sp. 100-5 ACE plasmid (Fig. 6). A particularly high coverage of 75% of the *Bosea* ACE plasmid was detected in the dataset from activated sludge microbial communities from an anaerobic bioreactor operated at Hunan University (Changsha, China). These findings indicate the worldwide distribution and genetic conservation of the ACE degradation genes clusters and the ACE plasmid.

## DISCUSSION

The artificial sweetener ACE was once considered recalcitrant in WWTPs [4] but recently its emerging biodegradability was reported [10, 12, 13]. The earliest indication for biodegradation based on ACE monitoring in wastewater was found in WWTPs located in Queensland, Australia [10] and Eriskirch, Germany [12], demonstrating removal efficiencies of about 90% in 2012 and 2013, respectively. Microbial community surveys highlighted the importance of *Alpha*- and *Gammaproteobacteria* for ACE degradation [13, 15]. In particular, representatives of the alphaproteobacterial genera *Bosea* and *Chelatococcus* capable of degrading ACE were isolated [14, 15]. The present study revealed the enzymatic and genetic background of ACE degradation for the first time and offers a glimpse of the evolutionary mechanisms driving the fast and worldwide spread of this novel xenobiotics degradation trait.

The comparison of the wildtype *Bosea* sp. 100-5 and its ACE degradation-defective mutant clearly revealed the involvement of a plasmid-borne gene cluster (ACE cluster 1) encoding an MBL-type hydrolase (BOSEA1005_40015) in ACE degradation. In line with this, the complete gene cluster including the flanking transposon-related CDS is highly conserved among all ACE-degrading strains analyzed in the present study. However, BOSEA1005_40015 appears to be unique and unprecedented, as the gene was not found in any other genomic context, and no closely related enzyme has been characterized thus far. In analogy to the distantly related hydrolases attacking phosphate ester bonds in nucleotides, the BOSEA1005_40015 enzyme might attack the sulfonyloxy group in ACE. The required translocation of the negatively charged substrate into the cell across the cytoplasmic membrane and against the membrane potential is likely enabled by the ABC-type transporter encoded in ACE cluster 1 (Fig. 3 and 4b). Finally, as the ACE hydrolysis product sulfamate formed in the cytoplasm is not used by ACE-degrading bacteria [14], an export system like the BOSEA1005_40018 protein is probably needed for the efficient removal of the waste product. Hence, together with BOSEA1005_40016 encoding an amidase, the gene cluster seems to be perfect for the two-step hydrolytic degradation of ACE via ANSA to acetoacetate and sulfamate. However, BOSEA1005_40016 was lost in *Bosea* sp. 100-5 Mut1 but protein extracts still hydrolyzed ANSA. Moreover, heterologous expression of another amidase gene (BOSEA1005_40030) in *E. coli* identified its function in ANSA hydrolysis and disproved the involvement of BOSEA1005_40016. Therefore, the genetic makeup of ACE cluster 1 implies that it originally evolved for the degradation of other substrate(s) sharing only some features with ACE.

Compounds structurally related to ACE that could be likewise attacked by the MBL-type sulfatase BOSEA1005_40015 are other natural or synthetic sulfamic acid derivatives [32, 33], in particular, O- and N-substituted sulfamates. In this context, Fig. S2a proposes a two-step enzymatic hydrolysis of ascamycin and other natural or synthetic aminoacyl sulfamate ribonulceoside antibiotics [34], involving sulfatase and amidase activities. Also sulfonamide antibiotics might be relevant, albeit having a deviating structure compared to sulfamates, as the sulfur atom of the sulfonyl group is directly linked to an aniline ring system (Fig. S2b). Although bacterial degradation seems to proceed mainly via ipso-hydroxylation leading to the decomposition of the resulting intermediates and the release of sulfite [35], alternatively, enzymatic hydrolysis by attacking the sulfur atom of, e.g., sulfacetamide, might lead to the release of aniline and an ANSA-related N-sulfonated acyl amide (Fig. S2b). In support, aniline as degradation product not compatible with the ipso-hydroxylation mechanism has been reported for *Pseudomonas psychrophila* HA-4 incubated with the sulfonamide antibiotic sulfamethoxazole [36]. Interestingly, the latter and other sulfonamides are partially removed from municipal wastewater concomitantly with ACE [11].

The ANSA amidase gene BOSEA1005_40030 is located in a second gene cluster on the ACE plasmid. In *Bosea* sp. 100-5, this cluster only consists of CDS for a transcriptional regulator (LysR) and the ANSA amidase. In *Chelatococcus* sp. 1g-11, additional genes encoding an anion import system and another TauE/SafE-like exporter are present (Figs. 5 and S3). In contrast to BOSEA1005_40015, sequences closely related to BOSEA1005_40030 can be found in genetic environments other than the ACE plasmid, e.g., WP_176954172.1 from *Paraburkholderia sartisoli* LMG 24000 showing 66% identity at 99% coverage. Interestingly, the gene clusters in strains 1g-11 and LMG 24000 share not only the amidase gene but also that encoding the anion import system (75% identity at 93% coverage) (Fig. S3). Additionally, a TauE/SafE system is present in strain LMG 24000, albeit not closely related to the ones encoded on the ACE plasmids. This constellation appears to have evolved for uptake of an extracellular amide that is hydrolyzed in the cytoplasm. *Bosea* sp. 100-5 lacks the TauE/SafE system, likely due to insertions of the IS5 family transposase and subsequent rearrangements, and the anion transport gene is only present as a pseudogene outside ACE cluster 2 (Fig. 3, Table 2). Hence, only the BOSEA1005_40030 amidase is needed for ANSA degradation but not the anion import system present in strain 1g-11.

Considering the unique and conserved gene arrangement, we propose that both gene clusters found on the ACE plasmids encode complete degradation pathways comprising substrate uptake, hydrolytic steps and export of a non-degradable product. The gene clusters seem to be highly evolved and perfectly coordinated for the degradation of yet unknown substrates. As outlined above, these substrates might be compounds structurally related to ACE and ANSA, likewise degradable by hydrolytic attack to organic intermediates that can be assimilated and dead end products to be removed from the cell.

Metabolic genes embedded in transposons and their location on a conjugative plasmid are typical features of bacterial pathways involved in degradation of xenobiotics, such as antibiotics and pesticides [37]. This enables the evolution and distribution of novel degradation pathways or other functional traits that facilitate niche adaptation. In case of ACE, mediated by the IS5 family transposase, ACE cluster 1 can easily transpose between different replicons (Fig. 5), which might have supported the recruitment of metabolic genes for ACE degradation. Probably, this development is not yet completed, as the ANSA amidase gene is found on the second plasmid-borne gene cluster. Further evolution might lead to the replacement of BOSEA1005_40016 by BOSEA1005_40030, resulting in a perfect ACE cluster.

From the biochemist’s perspective, ACE represents a challenging molecule combining features of a carboxylic acid amide and a sulfamic acid ester in an electron-rich ring system. Particularly, nucleophilic attack of the amide is difficult due to the localization of the negative charge mainly at the nitrogen atom [38]. Furthermore, even when used for productive degradation, growth yields on ACE would be low, as (i) its uptake likely depends on ATP hydrolysis and (ii) only the carbon skeleton could be used for assimilation and dissimilation (representing only 50% of the molecular weight). Hence, it can be concluded that nature lacked an appropriate degradation mechanism when ACE was introduced into the environment for the first time in the 1980s [2]. In agreement, the artificial sweetener was long time considered recalcitrant against enzymatic attack [4]. The unprecedented high removal efficiency monitored in WWTPs since 2012 [10] points to the evolution of an ACE degradation pathway within a 30-years period. However, considering its low concentration in wastewater (typically <100 μg L^−1^ in Europe, China and Australia [4, 8, 10, 12]), ACE is a rather poor carbon and energy source. Consequently, its utilization is likely not a sufficient driving force for the evolution of such a well-coordinated degradation pathway as depicted in Fig. 4b. Rather, co-metabolism with more dominant environmental chemicals can be expected [39]. In line with this, no earlier stage or any other evolution of ACE cluster 1 was traceable in genomes and metagenomes. However, older metagenome datasets might have insufficient sequencing depth to cover such rare genes. Moreover, MAGs are prone to miss variable genes of natural populations [40]. Accordingly, matches for the genes of the ACE clusters were only found in partially assembled metagenome and metatranscriptome data (Fig. 6, Table S2) but not in MAGs. Direct and systematic Sequence Read Archive (SRA) search would be needed for tracing gene fragments that are discarded by assembly algorithms.

The concomitant occurrence of identical gene clusters in Germany and China (Figs. 5 and 6) suggests that evolution and distribution of the individual clusters started much earlier than in the 2010s and was likely triggered by other natural or anthropogenic chemicals that exerted higher selective pressure than the non-toxic ACE. However, the last evolutionary step of combining the two gene clusters in one genome can be attributed to the recent bacterial adaptation to ACE as energy and carbon source. Due to the low ACE concentration in wastewater, this evolution might have occurred in rather oligotrophic environments, such as treatment wetlands [13, 41] and biofilms thriving in receiving waters. Further isolation studies and molecular surveys using the ACE and ANSA hydrolases as genetic markers could reveal how the ACE degradation trait has spread in aquatic environments worldwide.

## DATA AVAILABILITY

Supplementary text, figures and tables are provided in the Supplementary information. The genomes of the bacterial strains were deposited at NCBI under the accession numbers given in Table 1.

## ACKNOWLEDGEMENTS

We thank Ute Lohse (UFZ) for excellent technical assistance, particularly, in strain cultivation and genome sequencing. Cloud computing facilities used for the bioinformatics analyses were provided by the BMBF-funded de.NBI Cloud within the German Network for Bioinformatics Infrastructure (de.NBI) (031A537B, 031A533A, 031A538A, 031A533B, 031A535A, 031A537C, 031A534A, 031A532B).

## AUTHOR CONTRIBUTIONS

SK, TR, TRe, LA and CD conceived and directed the project. DP, MB, KL, CS, TR, YL and CA performed the experiments and were involved in method design as well as data analysis. MB, TR and SK wrote the manuscript with substantial input of all the authors.

