## Supplementary text, figures and tables for "Recently evolved combination of unique sulfatase and amidase genes enables bacterial degradation of the wastewater micropollutant acesulfame worldwide"

[#] Current affiliation: Institute of Human Genetics, University of Leipzig Medical Center, Leipzig, Germany

[‡] Current affiliation: Fraunhofer Institute for Interfacial Engineering and Biotechnology, Stuttgart, Germany

The Supplementary information comprises

**NCBI and JGI database search for BOSEA1005_40015 and other ACE plasmid genes……….……………3**

**Detailed description of the LC-MS/MS method for monitoring ANSA……………………..…..…………….3**

**Functional assignment for predicted proteins encoded in ACE cluster 1……….………………………….4**

**Functional assignment for predicted proteins encoded in ACE cluster 2……………………………..……..5**

**Table S1 Features of the bacterial genomes sequenced in this study.……………………….……………..……7**

**Table S2 NCBI and JGI metagenome and metatranscriptome**
**datasets that presented the ACE hydrolase gene…………………………………..……………..................8**

**Comparison of *Bosea* sp. 100-5 wildtype and Mut1 genomes (Fig. S1)…………….………………………9**

**Possible two-step enzymatic hydrolysis of ACE-related chemicals (Fig. S2)…………………………………..10**

**Comparison of gene clusters encoding the ANSA hydrolase (Fig. S3)……………………………..…………12**

**References…………………………………………………………………………………………….………………13**

### NCBI and JGI database search for BOSEA1005_40015 and other ACE plasmid genes

NCBI and JGI databases were searched using different strategies. In NCBI, we used blastp with both nr and env_nr databases as well as tblastn with the whole-genome shotgun contigs (wgs) database. The tblastn search was limited to the following metagenomes: activated sludge (taxid:942017), aquatic (taxid:1169740), aquifer (taxid:1704045), bioreactor (taxid:1076179), bioreactor sludge (taxid:412754), drinking water (taxid:2651591), freshwater (taxid:449393), groundwater (taxid:717931), lagoon (taxid:1763544), lake water (taxid:1647806), pond (taxid:1851193), sludge (taxid:1592332) and wastewater (taxid:527639). In JGI, blastp search (E-value < 1 e-50) was done in the metagenome datasets wastewater treatment plant (354) and wastewater (416), and in the metatranscriptome datasets wastewater treatment plant (43) and wastewater (20). Metagenome dataset bioreactor (798) and metatranscriptome dataset bioreactor (258) were also investigated after filtering study names containing “activated sludge” or “wastewater”.

### Detailed description of the LC-MS/MS method for monitoring ANSA

We used an Agilent 1260 Infinity II Series liquid chromatography (LC) system coupled to an AB Sciex QTRAP® 6500+ tandem mass spectrometer (MS/MS) equipped with a Turbo V™ ion source operated with electrospray ionization (ESI) in negative polarity. The enhanced product ion (EPI) spectrum of acetoacetamide-N-sulfonate (ANSA) shown in Fig. 2b was obtained at a scan rate of 10 000 Da $s^{-1}$ using dynamic fill time at a declustering potential of -20 V, an entrance potential of 10 V, a collision energy of -25 V and a collision energy spread of 15 V across the *m/z* range of 55 to 200 for structural elucidation. The ion source-dependent MS parameters, which were kept constant during the whole acquisition, were as follows: curtain gas (CUR): 30 psi; ion spray voltage (IS): -4 500 V; turbo spray temperature (TEM): 450 °C; nebulizer gas (GS1): 60 psi; heater gas (GS2): 60 psi; CAD gas: medium. Nitrogen was used as the curtain and collision gas.

The compound-dependent MS parameters (declustering potential (DP), entrance potential (EP), collision energy (CE) and collision cell exit potential (CXP)) were optimized by direct infusion of the analyte. We used multiple reaction monitoring (MRM) mode combined with the information dependent acquisition (IDA) criteria set at an intensity threshold of 5 000 cps, followed by an EPI as the dependent scan. The MRM transitions and optimized compound-dependent MS parameters for the analyte of interest are listed below.

| Analyte | Rt (min) | Precursor ion (*m/z*) | Product ions (*m/z*) | DP (V) | EP (V) | CE (V) | CXP (V) | Dwell time (ms) |
|---|---|---|---|---|---|---|---|---|
| ANSA | 1.19 | 179.9 | 95.8/79.8 | -20/-20 | -10/-10 | -20/-50 | -15/-16 | 40 |

All data were acquired and processed using Analyst 1.7.2 Software.

### Functional assignment for predicted proteins encoded in ACE cluster 1

**BOSEA1005_40011, _40012 and _40013:** ATP-binding cassette (ABC) import system with BOSEA1005_40011 encoding the ATP-binding protein, BOSEA1005_40012 the transmembrane subunit and BOSEA1005_40013 the periplasmic substrate-binding protein. Protein sequences of the importer are distantly related to uptake systems associated with the transport of small inorganic or organic anions. However, only very low identity to thus far studied proteins can be found, e.g., the periplasmic substrate-binding protein shows 27% sequence identity at 18% query coverage to TauA from *E. coli* (UniProt ID Q47537), which was reported to bind preferentially the short-chain alkanesulfonate taurine (Qu, ElOmari et al. 2019).

**BOSEA1005_40015:** Metallo beta-lactamase (MBL)-type ACE hydrolase, 283 aa and with a predicted molecular weight of 34.9 kDa. Conserved residues for coordination of two divalent metal ions per subunit are H61, H63, D65, H66, H172 and H251. In the protein data bank (PDB), substantial sequence coverage with the BOSEA1005_40015 protein is only provided by Apyc1 from *Bacillus* phage BSP38 (PDB ID 7T28) showing 26% identity (90% query coverage). The phage protein catalyzes the hydrolysis of cyclic mononucleotides involved in the antiphage defense system of its host. Its di-metal center is occupied by $Zn^{2+}$ ions (Hobbs, Wein et al. 2022). Other distantly related hydrolases (only <30% identical residues at >80% query coverage) are often annotated as the short form of ribonuclease Z (consisting of 300 to 400 aa) that removes extra 3' nucleotides from tRNA precursors (Takaku, Minagawa et al. 2004). From the specific metal dependence of ACE degradation observed in *Bosea* sp. 100-5 (Fig. 2d), it can be concluded that the reaction center of the active MBL-type hydrolase is exclusively occupied by two $Mn^{2+}$ ions. This is somewhat surprising, as MBL-type enzymes may be able to coordinate other divalent metal ions when the favored species is not available (Hu, Spadafora et al. 2009). Presumably, in the case of the ACE hydrolase, the substitution with other metals than $Mn^{2+}$ results in a substantial reduction in activity. This might be explained by the ACE structure that obviously requires the full activity of the enzyme, as a nucleophilic attack towards the sulfur or C3 atom (Fig. 1) is quite challenging due to the electron-richness of the ring system.

**BOSEA1005_40016:** Amidase bearing the amidase signature sequence, 465 aa and with a molecular weight of 49.3 kDa. Residues of the catalytic Ser Ser Lys triad for attacking amide groups are K81, S156 and S180. Related sequences (about 60% identity at 96% query coverage) can be found in gene environments other than the ACE degradation gene cluster, e.g., WP_013973238.1 (nicotine-degrading *Pseudomonas putida* S16) and WP_244145082.1 (root nodule bacterium *Paraburkholderia tuberum* LMG 21444). However, as these matches lack biochemical characterization, no conclusion on substrate preferences can be drawn.

**BOSEA1005_40018:** Belongs to the TauE/SafE family of transmembrane proteins involved in the export of small anions, such as sulfite, sulfoacetate and 3-sulfolactate (Weinitschke, Denger et al. 2007, Mayer, Denger et al. 2012). The TauE/SafE system is not well characterized and the amino acid sequence is poorly conserved among representatives of this protein family, e.g., BOSEA1005_40018 does not show significant similarity (blastp) with WP_011617520.1 from *Cupriavidus necator* H16.

### Functional assignment for predicted proteins encoded in ACE cluster 2

**CHELA1g11_60039:** 450 aa protein, permease system that can likely operate as a chemiosmotic transporter for the uptake of small anions. Distantly related to a putative short-chain fatty acid transporter found in *E. coli* (UniProt ID P76460; 34% identity at 94% query coverage). In *Bosea* sp. 100-5 and *Bosea* sp. 3-1B, the homologous gene is disrupted/incomplete and not part of the gene cluster (Fig. 3). See also Fig. S3 (P. 12) for comparison.

**CHELA1g11_60040 / BOSEA31B_30045:** Another TauE/SafE export system (see ACE cluster 1, BOSEA1005_40018) involved in excretion of small anions. The CHELA1g11_60040 / BOSEA31B_30045 protein shows only low similarity to BOSEA1005_40018 (44% identity at 10% query coverage, blastp). See also Fig. S3 (P. 12) for comparison.

**BOSEA1005_40029:** The gene product (311 aa) is predicted to be a helix-turn-helix domain bearing transcriptional regulator of the LysR family. In the UniProt database, one of the closest matches (30% identity 78% query coverage) is AlsR from *Bacillus subtilis* 168 (ID Q04778), which is involved in regulating the expression of acetoin synthesis genes (Renna, Najimudin et al. 1993). Due to the highly conserved co-occurrence of BOSEA1005_40029 and BOSEA1005_40030 sequences in ACE cluster 2 found in ACE-degrading strains, the LysR family protein might be involved in the regulation of the expression of the ANSA amidase gene.

**BOSEA1005_40030:** ANSA amidase bearing the amidase signature sequence. The total protein possesses 472 aa and a molecular weight of 50.5 kDa. Residues of the catalytic Ser Ser Lys triad for attacking amide groups are K83, S158 and S182. The closest match of a characterized enzyme is an amidase from *Rhodococcus erythropolis* TA37 (UniProt ID K9NBS6) showing 40% identity at 95% query coverage. This amidase from strain TA37 is specific for short-chain N-substituted amides (Lavrov, Zalunin et al. 2010). Likewise, BOSEA1005_40030 is only distantly related to BOSEA1005_40016 (amidase 1) of the sulfatase gene cluster (sharing only 39% identity residues at 90% query coverage). As discussed in the manuscript, a sequence very similar to BOSEA1005_40030 can be found in *Paraburkholderia sartisoli* LMG 24000. See also Fig. S3 (P. 12) for comparison. However, the latter strain does not contain the ACE hydrolase (BOSEA1005_40015).

**Table S1 Features of the bacterial genomes sequenced in this study.**

| Strain | ***Bosea* sp. 100-5** | ***Bosea* sp. 100-5 Mut1** | ***Bosea* sp. 3-1B** | ***Chelatococcus* sp. 1g-2** | ***Chelatococcus* sp. 1g_11** | ***Chelatococcus asaccharovorans* WSA4-1** | ***Chelatococcus* sp. WSC3-1** | ***Chelatococcus asaccharovorans* WSD1-1** | ***Chelatococcus* sp. WSG2-a** |
|---|---|---|---|---|---|---|---|---|---|
| **Accession number** | GCA_930633465 | GCA_946047605 | GCA_930633495 | GCA_930633525 | GCA_930633505 | GCA_930633515 | GCA_930633455 | GCA_930633485 | GCA_930633475 |
| **Genome size (bp)** | 5 939 164 | 5 829 812 | 5 944 673 | 7 159 619 | 7 252 269 | 7 194 471 | 6 967 327 | 7 195 039 | 7 037 809 |
| **Number of contigs** | 4 | 68 | 3 | 6 | 8 | 6 | 5 | 6 | 5 |
| **GC content (%)** | 65.78 | 65.85 | 65.78 | 62.81 | 62.76 | 64.14 | 63.01 | 64.15 | 62.98 |
| **Total number of CDS** | 6 101 | 6 014 | 6 086 | 7 265 | 7 381 | 7 193 | 6 914 | 7 190 | 7 024 |
| **Number of tRNA genes** | 63 | 59 | 63 | 55 | 55 | 51 | 54 | 51 | 55 |
| **Number of rRNA genes (5S, 16S, 23S)** | 6 (2, 2, 2) | 3 (1,1,1) | 6 (2, 2, 2) | 9 (3, 3, 3) | 9 (3, 3, 3) | 6 (2, 2, 2) | 9 (3, 3, 3) | 6 (2, 2, 2) | 9 (3, 3, 3) |
| **Number of miscella-neous RNA genes** | 22 | 22 | 22 | 37 | 39 | 41 | 34 | 40 | 35 |
| **Pseudo-genes** | 10 | 6 | 7 | 2 | 2 | 0 | 0 | 0 | 0 |
| **CheckM Complete-ness (%)** | 99.05 | 98.9 | 99.05 | 99.68 | 99.68 | 100 | 99.37 | 100 | 99.37 |
| **CheckM Contamina-tion (%)** | 2.92 | 2.92 | 2.92 | 1.35 | 1.35 | 1.41 | 1.61 | 1.41 | 1.3 |
| **Taxonomy (GTDB-tk)** | *Bosea* | *Bosea* | *Bosea* | *Chelatococcus* | *Chelatococcus* | *Chelatococcus asaccharo-vorans* | *Chelatococcus* | *Chelatococcus asaccharo-vorans* | *Chelatococcus* |

**Table S2 NCBI and JGI metagenome and metatranscriptome datasets that presented the ACE hydrolase gene.** The BOSEA1005_40015 sequence was used as query (protein with 283 aa). Datasets were obtained from activated sludge samples of various sources as indicated. Numbering as given in Fig. 6.

| Number | Source | Accession | Description | Query coordinates | Query coverage (%) | Identity (%) | E value | Sampling date | Geographic location |
|---|---|---|---|---|---|---|---|---|---|
| **NCBI database** | | | | | | | | | |
| **6** | Metagenome | WBZQ 01256123.1 | nitrification reactor of WWTP[a] location 7 | 1…150 | 53 | 99 | 4E-088 | Sept. 2017 | Virginia, USA |
| **6** | Metagenome | WBZQ 01256123.1 | nitrification reactor of WWTP[a] location 7 | 155…283 | 46 | 99 | 2E-073 | Sept. 2017 | Virginia, USA |
| **JGI database** | | | | | | | | | |
| **3** | Metagenome | 3300009873 | WWTP, Wenshan plant | 1…238 | 84 | 100 | 2E-178 | Dec. 2015 | Taichung, Taiwan |
| **2** | Metagenome | 3300009870 | WWTP, Linkou plant | 1…144 | 51 | 100 | 1E-103 | Dec. 2015 | New Taipei, Taiwan |
| **5** | Metatranscriptome | 3300007323 | Klosterneuburg WWTP, MT KNB_C2_LD | 1…150 | 53 | 99 | 4E-109 | March 2015 | Klosterneuburg, Austria |
| **5** | Metatranscriptome | 3300006644 | Klosterneuburg WWTP, MT KNB_A2_L | 1…109 | 39 | 99 | 1E-75 | March 2015 | Klosterneuburg, Austria |
| **5** | Metatranscriptome | 3300007595 | Klosterneuburg WWTP, MT KNB_T0_6L | 177…255 | 28 | 100 | 9E-52 | March 2015 | Klosterneuburg, Austria |
| **4** | Metatranscriptome | 3300028647 | WWTP Weurt | 1…283 | 100 | 99 | 0E+00 | Aug. 2017 | Nijmegen, Netherlands |
| **1** | Metagenome | 3300022303 | anaerobic bioreactor, Hunan University, 11 | 1…283 | 100 | 99 | 0E+00 | (2017)[b] | Changsha, China |
| **1** | Metagenome | 3300022241 | anaerobic bioreactor, Hunan University, 6 | 62…283 | 78 | 99 | 1E-164 | (2017)[b] | Changsha, China |
| **1** | Metagenome | 3300022302 | anaerobic bioreactor, Hunan University, 3 | 87…283 | 70 | 99 | 1E-143 | (2017)[b] | Changsha, China |
| **1** | Metagenome | 3300022302 | anaerobic bioreactor, Hunan University, 3 | 1…66 | 23 | 100 | 8E-40 | (2017)[b] | Changsha, China |
| **1** | Metagenome | 3300022215 | anaerobic bioreactor, Hunan University, 7 | 1…152 | 54 | 99 | 3E-110 | (2017)[b] | Changsha, China |
| **1** | Metagenome | 3300022238 | anaerobic bioreactor, Hunan University, 4 | 22…144 | 43 | 98 | 1E-86 | (2017)[b] | Changsha, China |
| **1** | Metagenome | 3300022291 | anaerobic bioreactor, Hunan University, 5 | 47…166 | 42 | 99 | 4E-85 | (2017)[b] | Changsha, China |
| **1** | Metagenome | 3300022291 | anaerobic bioreactor, Hunan University, 5 | 232…283 | 18 | 98 | 1E-26 | (2017)[b] | Changsha, China |

[a]WWTP – wastewater treatment plant

[b]sampling date not available, database entry created Dec. 2017

#### Comparison of *Bosea* sp. 100-5 wildtype and Mut1 genomes

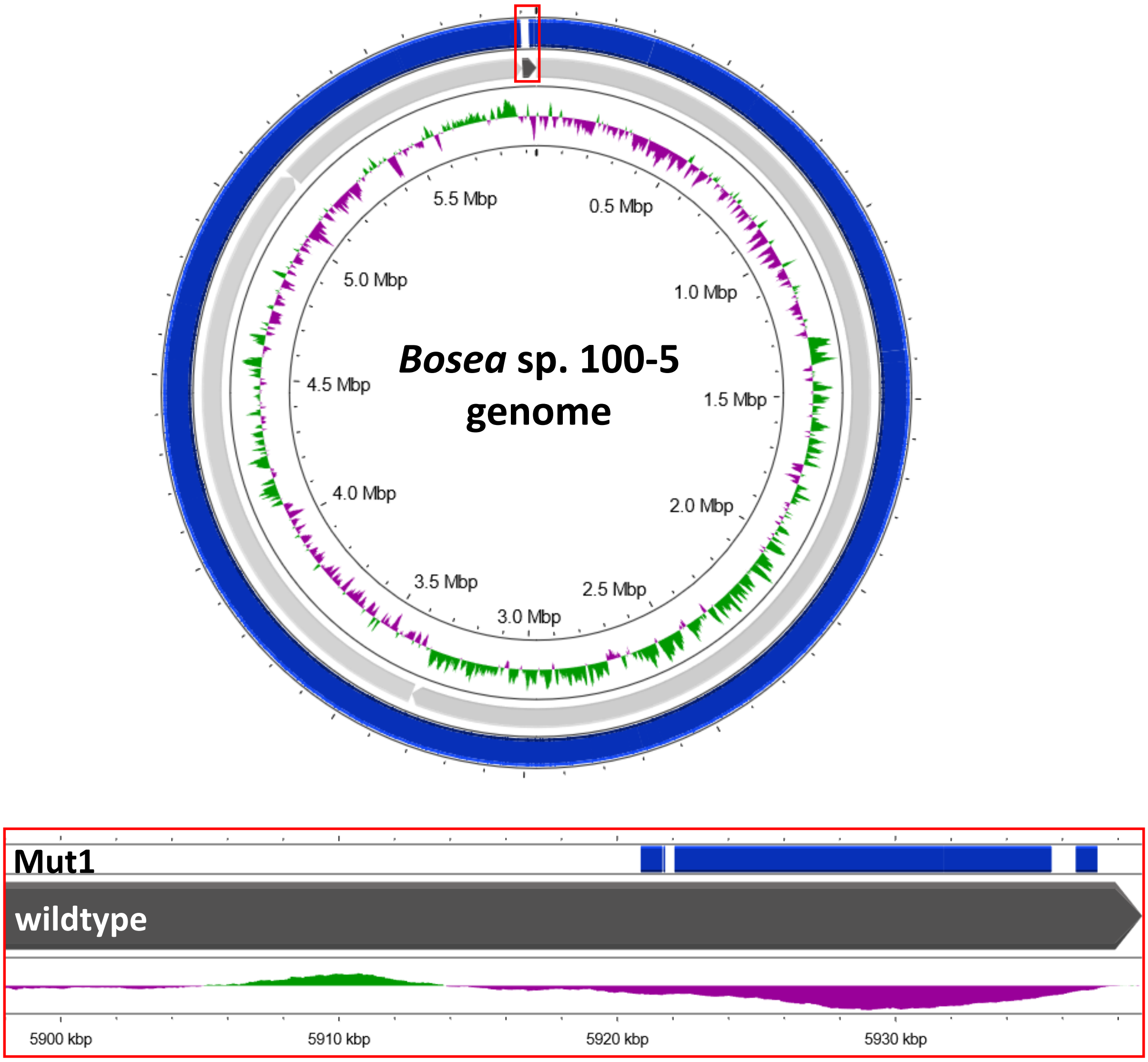


**Fig. S1 Alignment of the *Bosea* sp. 100-5 Mut1 genome (blue) against the *Bosea* sp. 100-5 genome (grey).** The 41-kb zoom (red framed box) shows the gaps found in the Mut1 genome, amounting to a total deletion of 26.3 kb. The figure was generated with Proksee.

### Possible two-step enzymatic hydrolysis of ACE-related chemicals

Among chemicals structurally related to ACE are, e.g., the anticonvulsant and antiepileptic drug topiramate (Privitera 1997) and antiviral nucleotide sulfamates (Winum, Scozzafava et al. 2005). A hydrolytic attack at the sulfur atom of these O-substituted sulfamates may release sulfamic acid/the sulfamate anion. However, regarding the subsequent amidase reaction, also an N-derivatization of the parental compound with a carbonyl carbon would be a prerequisite for the evolution of a two-step hydrolase pathway for ACE degradation. Furthermore, sulfamic acid derivatives used as pharmaceuticals can only cause micropollutions in wastewater, thus, being insignificant growth substrates as well. Rather, organic sulfamates directly affecting bacterial growth could explain the evolution of the ACE gene clusters. In this context, ascamycin and other natural or synthetic aminoacyl sulfamate ribonulceosides acting as inhibitors of aminoacyl-tRNA synthetases might play a role. Although enzymatic hydrolysis in *Xanthomonas campestris* seems to affect initially the amide and is reported to be catalyzed by a proline iminopeptidase (UniProt ID P52279) not related to amidase signature sequence hydrolases (Sudo, Shinohara et al. 1996), the combination of a BOSEA1005_40015-like sulfatase with BOSEA1005_40016- or 40030-like amidases in other bacteria might neutralize the antibiotic activity and release sulfamic acid/sulfamate as one reaction product (Fig. S2a).

Likewise, sulfonamide antibiotics might be relevant, as these compounds have already been applied for many decades (Henry 1943, Seydel 1968) and resistance against them can be strongly selective. As shown in Fig. S2b, depending on the sulfonamide structure, a sulfatase-catalyzed attack might lead to formation of an amide-N-sulfonate intermediate that could be further hydrolyzed by an amidase to the corresponding carboxylic and sulfamic acid ions.

Biochemical characterization of the BOSEA1005_40015, _40016 and _40030 enzymes by elucidating structures and substrate specificity may be worthwhile and could extend our knowledge on sulfamate and sulfonamide degradation mechanisms. Currently, however, the origin of the individual ACE gene clusters and the evolutionary drivers that established the ACE pathway specifically in alphaproteobacterial genera remains elusive.

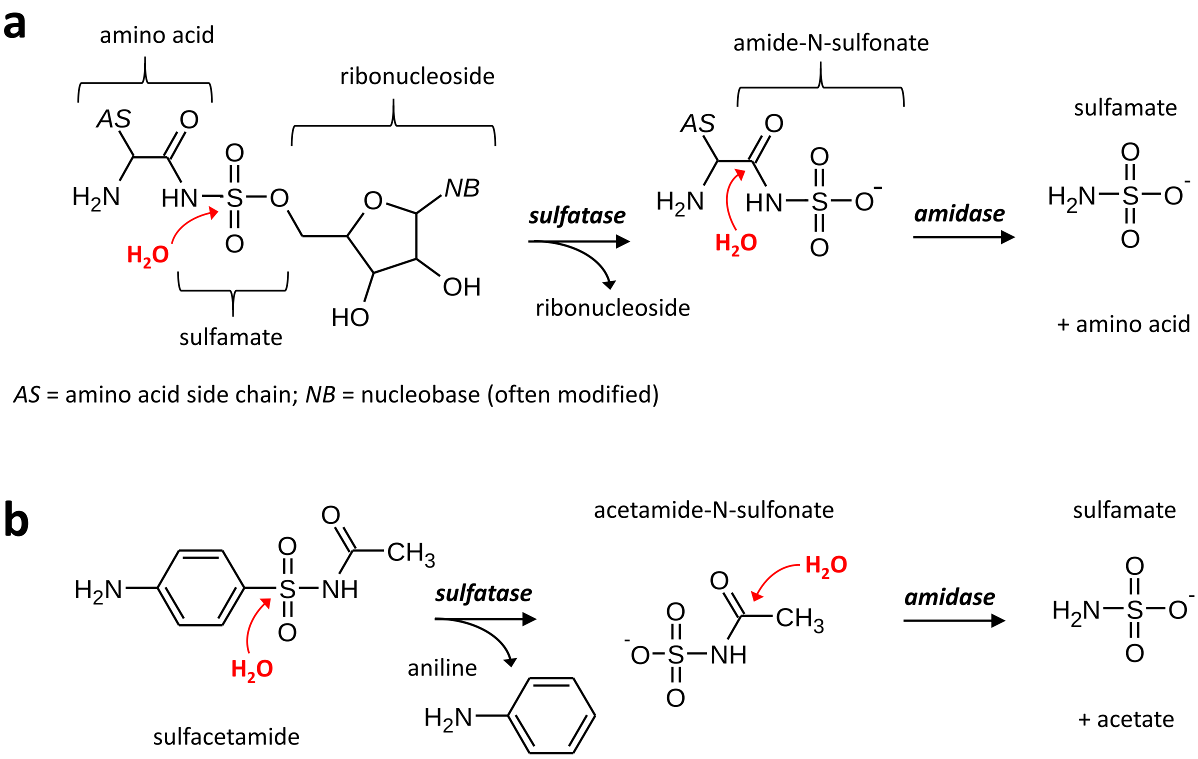


**Fig. S2 Hypothetical pathways for the two-step hydrolysis of ACE-related chemicals catalyzed by BOSEA40015_40015-like sulfatases and BOSEA1005_40016- or 40030-like amidases. a** Hydrolysis of aminoacyl sulfamate ribonucleotide antibiotics could proceed via an amide-N-sulfonate as intermediate and could form the sulfamate ion as end product. **b** Likewise, the sulfonamide antibiotic sulfacetamide might be degraded via acetamide-N-sulfonate.

#### Comparison of gene clusters encoding the ANSA hydrolase

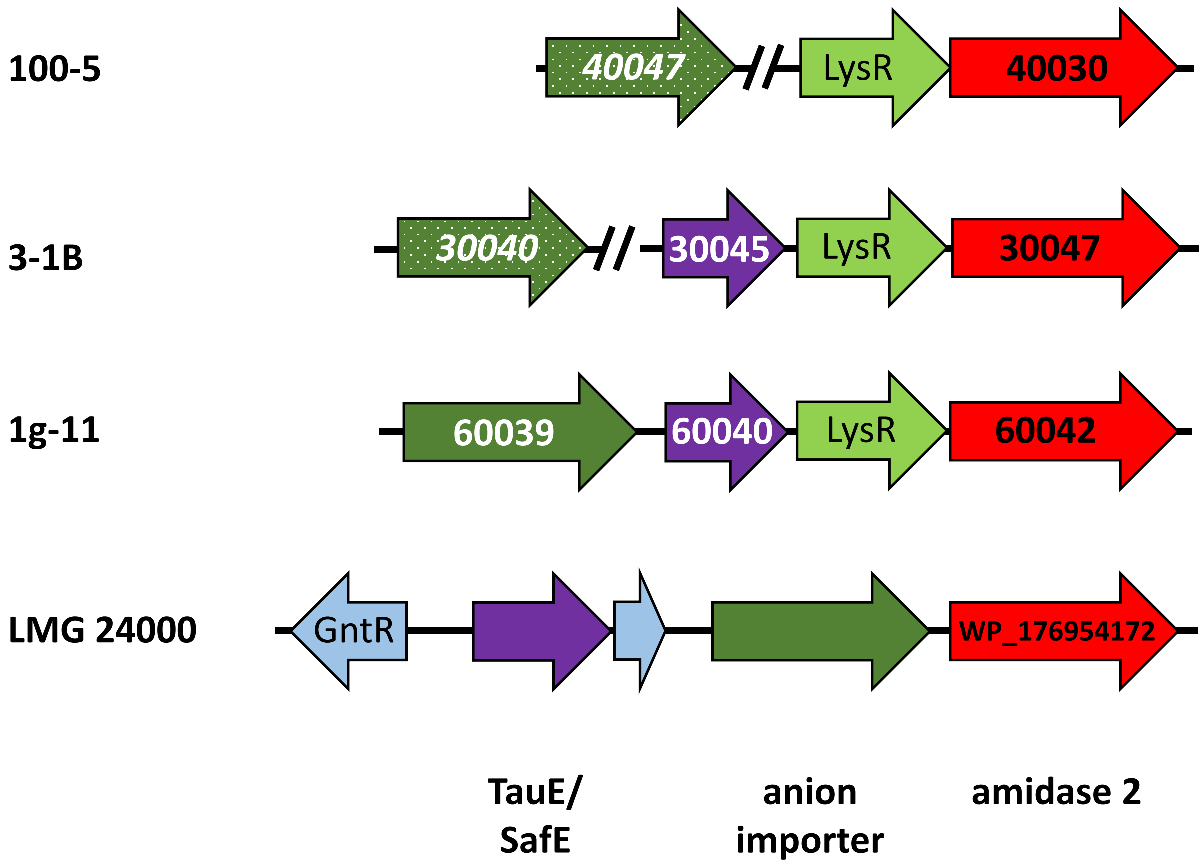


**Fig. S3 Comparison of gene clusters bearing the gene for the ANSA-hydrolyzing enzyme (amidase 2).** In *Bosea* sp. 100-5, *Bosea* sp. 3-1B and *Chelatococcus* sp. 1g-11, the cluster is found on the ACE plasmid, whereas it is located in *Paraburkholderia sartisoli* LMG 24000 on a contig of 454 572 bp (NZ_FNRQ01000004.1) that is likely part of the chromosome. Sequences of the amidase, the TauE/SafE exporter (absent in *Bosea* sp. 100-5) and the LysR-like transcriptional regulator are identical in the ACE degraders. The anion importer is only complete in strain 1g-11 (450 aa). In the *Bosea* strains, it can be interpreted as a pseudogene (BOSEA1005_40047, BOSEA31B_30040) located outside the gene cluster and resulting in a protein with missing N- and C-termini. The CDS start has been disrupted by the IS5 family transposase (Fig. 3). In total, only about 330 aa can still be aligned to CHELA1g11_60039 and related transporters. In contrast, the importer is conserved in strain LMG 24000 (sharing 75% identity at 93% coverage with CHELA1g11_60039). Likewise, the amidase of this strain is closely related to the ANSA amidase (66% identity at 99% coverage). The TauE/SafE system, however, does not show significant match (blastp) on sequence level to the protein encoded in the amidase gene cluster of the ACE degraders.
